## Supplementary material for "Pharmacological inhibition of astrocytic transglutaminase 2 facilitates the expression of a neurosupportive astrocyte reactive phenotype in association with increased histone acetylation": Figures S1-S4, table S5

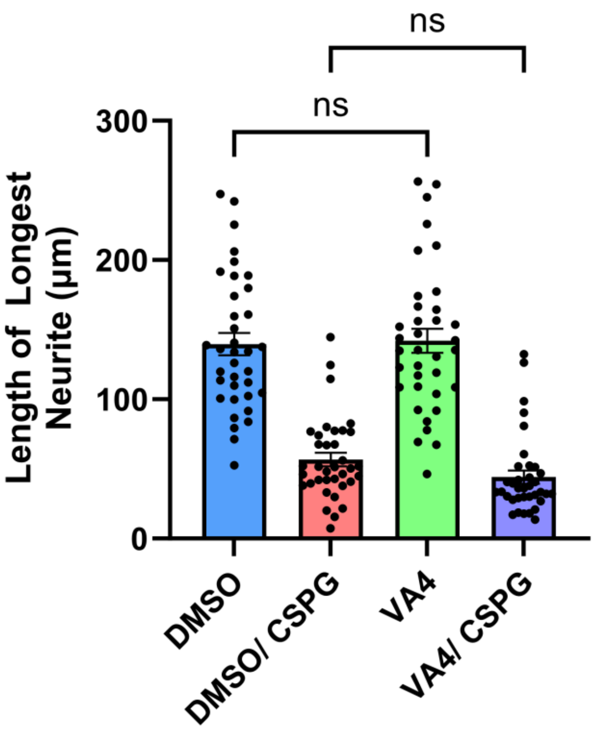


**Figure S1.** VA4 treatment of neurons does not affect neurite outgrowth on poly D lysine (PDL) or chondroitin sulfate proteoglycan (CSPG) matrices. Neurons were directly treated with DMSO (control) or 10 µM VA4 for 4 days prior to quantification of neurite length on PDL or CSPG matrices. Shown as Mean and SEM (n=36 neurons per condition from 1 biological replicate, Two-way ANOVA).

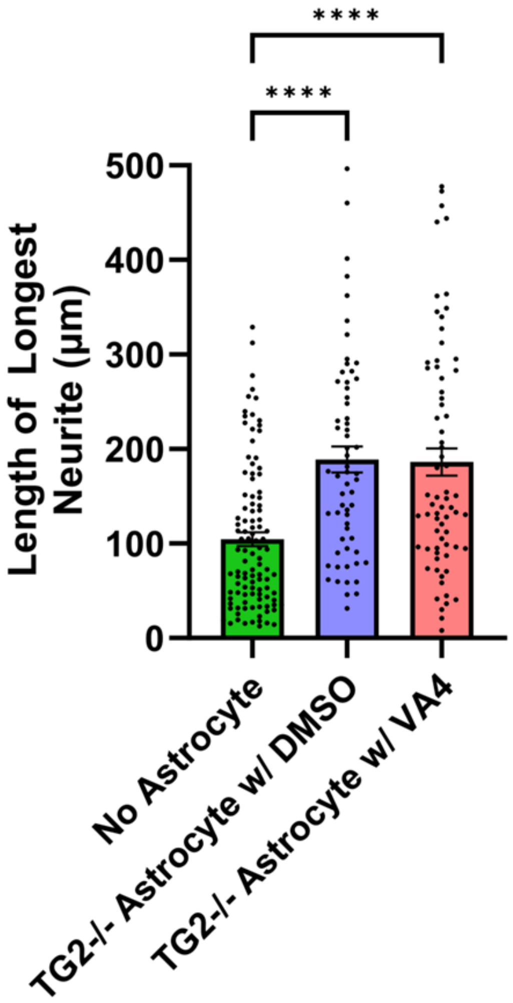


**Figure S2.** VA4 treatment of TG2-/- astrocytes does not further facilitate their ability to promote neurite outgrowth on chondroitin sulfate proteoglycan (CSPG) matrices. Astrocytes were treated with DMSO (control) or 10 µM VA4 for 4 days prior to pairing with neurons on CSPG matrices. Shown as Mean and SEM (n=61-106 neurons per condition from 1 biological replicate, Kruskal-Wallis Test).

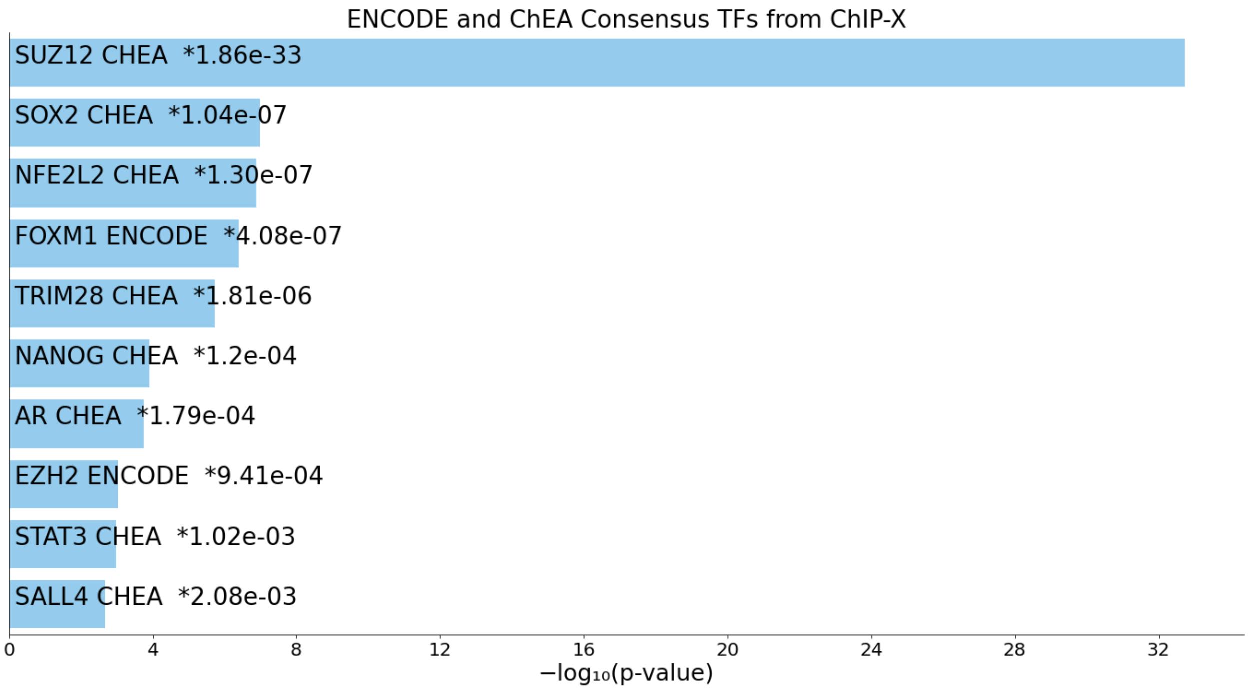


**Figure S3.** Enrichr transcription factor gene ontology of differentially expressed genes in TG2-/- astrocytes from previous RNA-seq data (Monteagudo et al., 2018) shows a strong enrichment in genes associated with PRC2 repressive chromatin modifying complex components (SUZ12 and EZH2) .

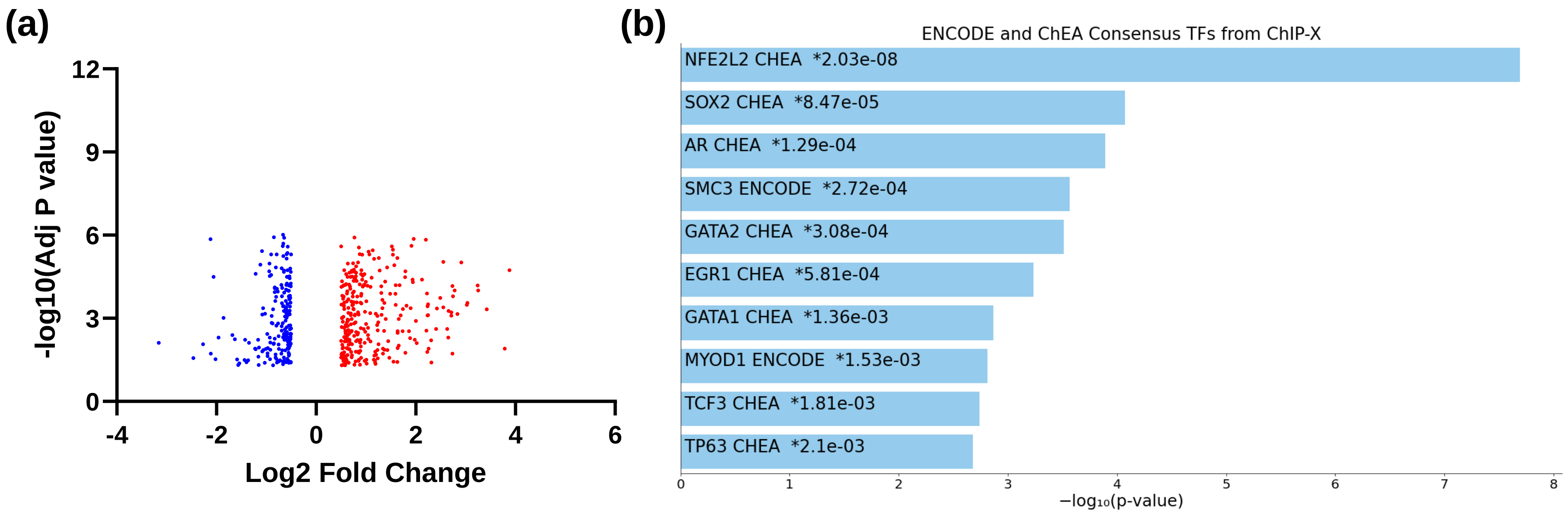


**Figure S4.** Replicate proteomic analysis of TG2-/- and WT astrocytes using data independent acquisition (DIA). Fold change represents the protein abundance ratio in TG2-/- astrocytes to WT astrocytes. Adj P value was calculated using Benjamini-Hochberg correction for multiple comparisons. Each experimental group contains two biological replicates with 3 technical replicates each for a total of 6 replicates per group.

**Table S5.** Detailed report of statistical methods used for each figure.

| | Normality Test p-value | # Outliers Removed | Test | Sided | Multiple Comparison Correction? | DF | Test Statistic | p-value |
|---|---|---|---|---|---|---|---|---|
| **Figure 1c** | p=0.1719, p=0.0911, p=0.0008 | 3, 0, 0 | Kruskal Wallis Test | N/A | Dunn's multiple comparisons test | 3, 300 | Z=5.814, Z=9.621, Z=3.251 | p<0.0001, p<0.0001, p=0.0034 |
| **Figure 2c** | p=0.3859 | 0, 0 | Unpaired T test | two | N/A | 7 | t=13.61 | p<0.0001 |
| **Figure 3b** | p=0.3387, p=0.2143 | 0, 1 | Unpaired T test | two | N/A | 11 | t=5.312 | p=0.0002 |
| **Figure 4b** | p=0.0917, p=0.0514 | 0, 0 | Unpaired T test | two | N/A | 3 | t=7.528 | p=0.0049 |
| **Figure S1** | p=0.2708, p=0.0103, p=0.1961, p=0.0017 | 0, 0, 0, 3 | Two-way ANOVA | one | Šídák's multiple comparisons test | 35, 3, 102 | F (35, 102) =1.191 F (3, 102) = 66.88 | p=0.2473 p<0.0001 |
| **Figure S2** | p=0.0001 p=0.0123 p=0.0004 | 2, 0, 0 | Kruskal Wallis Test | N/A | Dunn's multiple comparisons test | 3, 234 | Z=5.277 Z=4.909 Z=0.5256 | p<0.0001 p<0.0001 p>0.9999 |
